# Optimized Workflow for Enrichment and Identification of Biotinylated Peptides using Tamavidin 2-REV for BioID and Cell Surface Proteomics

**DOI:** 10.1101/2022.03.04.483072

**Authors:** Kohei Nishino, Harunori Yoshikawa, Kou Motani, Hidetaka Kosako

## Abstract

Chemical or enzymatic biotinylation of proteins is widely used in various studies, and proximity-dependent biotinylation coupled to mass spectrometry is a powerful approach for analyzing protein–protein interactions in living cells. We recently developed a simple method to enrich biotinylated peptides using Tamavidin 2-REV, an engineered avidin-like protein with reversible biotin-binding capability. However, the low abundance of protein biotinylation in cells required large amounts of cellular proteins to detect enough biotinylated peptides. Here we optimized the workflow for efficient enrichment and identification of biotinylated peptides. The most efficient recovery was achieved by heat inactivation of trypsin, prewashing Tamavidin 2-REV beads, clean-up of biotin solution, mock elution, and the optimal temperature and salt concentration for elution. Using the optimized workflow, over 2-fold more biotinylated peptides were identified with higher purity from RAW264.7 macrophages expressing TurboID-fused STING. In addition, sequential digestion with Glu-C and trypsin led to the identification of biotinylation sites that were not identified by trypsin digestion alone. Furthermore, the combination of this workflow with TMT labeling enabled large-scale quantification of cell surface proteome changes upon EGF stimulation. This workflow would be useful not only for BioID and cell surface proteomics but also for various other applications based on protein biotinylation.

## INTRODUCTION

Chemical or enzymatic biotinylation of proteins is widely used in various studies including affinity purification, enzyme-linked immunosorbent assay, Western blotting, cell staining, cell surface labeling, and identification of post-translational modifications (1, 2). Recently, proximity-dependent biotinylation approaches such as BioID and APEX have emerged as powerful techniques to identify protein–protein interactions and subcellular proteomes in living cells and organisms (3–5). Biotinylated proteins can be captured by streptavidin beads and identified by mass spectrometry after on-bead digestion or digestion of eluted proteins (6–10). However, the extremely high affinity of streptavidin for biotin makes it difficult to efficiently and specifically elute biotinylated proteins and remove contaminants of non-biotinylated proteins. While the direct identification of biotinylated peptides can eliminate these contaminants and provide further information about the topology of membrane proteins and the sites of post-translational modifications (11–14), biotinylated peptides are rarely detected after pull-down with streptavidin beads.

To efficiently enrich and identify biotinylated peptides, two groups have developed a revolutionary method using anti-biotin antibody beads, from which biotinylated peptides can be efficiently but non-specifically eluted with strong acid and high concentration of organic solvent (15, 16). Very recently, we developed a simple method to specifically enrich biotinylated peptides using Tamavidin 2-REV beads (17). Tamavidin 2-REV is an engineered version of Tamavidin 2 that acquired reversible biotin-binding capability by mutating S36 that forms a hydrogen bond with biotin (17, 18). Because biotinylated peptides can be mildly and specifically eluted from Tamavidin 2-REV beads by adding excess free biotin, the enrichment efficiency was higher than that of anti-biotin antibody beads. However, interference ions derived from our enrichment process prevented sensitive detection of biotinylated peptides. Furthermore, the low abundance of protein biotinylation in BioID-expressing cells required large amounts of cellular proteins to identify enough biotinylated peptides by mass spectrometry (17, 19). Therefore, it is important to optimize the enrichment method for biotinylated peptides, as various enrichment methods for phosphorylated peptides have been extensively optimized so far.

Here, we optimized each step of the workflow for efficient enrichment and identification of biotinylated peptides using Tamavidin 2-REV. Consequently, approximately 1,400 biotinylated peptides were identified from 300 μg proteins of RAW264.7 macrophages expressing TurboID-fused STING (stimulator of interferon genes). This number of identified biotinylated peptides was more than twice as many as before optimization, and approximately 95% of all identified peptides were biotinylated. We also found that sequential digestion with Glu-C and trypsin led to the identification of biotinylation sites that were not identified by trypsin digestion alone. Furthermore, the combination of this workflow with TMT labeling enabled large-scale quantification of biotinylated peptides from EGF-stimulated and then surface-biotinylated HeLa cells. This optimized workflow using Tamavidin 2-REV would be useful for BioID and cell surface proteome profiling as well as for various other applications based on protein biotinylation.

## MATERIALS AND METHODS

### Reagents

MagCapture HP Tamavidin 2-REV magnetic beads and lysyl endopeptidase (Lys-C) were purchased from FUJIFILM Wako Pure Chemical (Osaka, Japan). DMEM high glucose, penicillin-streptomycin solution, and D-biotin were from Nacalai Tesque (Kyoto, Japan). Fetal bovine serum (FBS) and Pefabloc SC were from Sigma-Aldrich (St. Louis, MO). EGF was from PeproTech (Rocky Hill, NJ). EZ-Link Sulfo-NHS-Biotin, BCA protein assay kit, trypsin (MS grade), and endoproteinase Glu-C (MS grade) were from Thermo Fisher Scientific (Waltham, MA). RapiGest SF was from Waters (Milford, MA). All other standard laboratory reagents were obtained from Nacalai Tesque, Sigma-Aldrich, or FUJIFILM Wako Pure Chemical.

### Cell Culture

STING-knockout RAW264.7 murine macrophages stably expressing TurboID-STING fusion protein (17, 20) and HeLa cells (21) were maintained in DMEM supplemented with 10% FBS, penicillin (100 U/mL), and streptomycin (0.1 mg/mL) at 37 °C in 5% CO_2_.

### EGF Stimulation and Cell Surface Biotin Labeling

Cell surface biotin labeling was performed according to previous reports with slight modifications (22, 23). HeLa cells in 6-cm dishes were treated with 100 ng/mL EGF for 0, 5, 10, or 30 min after starvation for 18 h in serum-free DMEM, washed twice with ice-cold PBS, and then incubated with EZ-Link Sulfo-NHS-Biotin (0.25 mg/mL in PBS) for 30 min at 4 °C with gentle agitation. The cells were then incubated with 0.1 M glycine in PBS for 10 min at 4 °C with gentle agitation. After washing with ice-cold TBS1 (20 mM Tris-HCl, pH 7.5, and 150 mM NaCl), the cells were scraped and collected by centrifugation at 4 °C.

### Protein Extraction and Enzymatic Digestion of BioID Samples

STING-knockout RAW264.7 cells stably expressing TurboID-STING were seeded in 6-cm dishes and cultured overnight. The cells at 80% confluence were incubated with biotin (500 μM final concentration) for 10 min, washed with ice-cold PBS, scraped on ice, collected by centrifugation at 4 °C, and lysed in 200 μL of guanidine buffer (6 M guanidine-HCl and 100 mM HEPES-NaOH, pH 7.5) containing 10 mM Tris (2-carboxyethyl) phosphine (TCEP) and 40 mM chloroacetamide (CAA). The lysates were heated at 95°C for 10 min and then sonicated at 4°C for 150 s (5 × 30 s pulses with 30 s intervals) at high power using a Bioruptor II (Cosmo Bio, Tokyo, Japan). Protein concentrations were quantified using a BCA assay kit. Then, 300 μg of proteins were purified by methanol–chloroform precipitation and resuspended in 200 μL of 0.1% RapiGest SF in 50 mM triethylammonium bicarbonate or 100 μL of PTS buffer [100 mM Tris-HCl, pH 8.0, 12 mM SDC, and 12 mM SLS (24–26)], followed by sonication and heating at 95°C for 10 min. The protein solution in PTS buffer was then diluted 5-fold with 100 mM Tris-HCl, pH8.0. The proteins were digested with trypsin (1:100) at 37 °C overnight, or with Lys-C (1:100) or Glu-C (1:100) at 37 °C for 4 h followed by digestion with trypsin (1:100) at 37 °C overnight.

### Original Workflow for Enrichment of Biotinylated Peptides

A 15-μL slurry of MagCapture HP Tamavidin 2-REV magnetic beads per sample was washed three times with TBS2 (50 mM Tris-HCl, pH 7.5, and 150 mM NaCl). The digested peptides containing 0.1% RapiGest SF were diluted 5-fold with TBS2 and incubated with the Tamavidin 2-REV beads in the presence of 1 mg/mL Pefabloc SC for 3 h at 4 °C. After washing five times with TBS2, biotinylated peptides were eluted with 100 μL of 1 mM biotin in TBS2 for 15 min at 95 °C twice. The combined eluates were desalted using GL-Tip SDB (GL Sciences, Tokyo, Japan), evaporated in a SpeedVac concentrator, and redissolved in 0.1% TFA and 3% acetonitrile (ACN).

### Optimized Workflow for Enrichment of Biotinylated Peptides

A 15-μL slurry of the Tamavidin 2-REV beads per sample was washed twice with 10% ACN and then three times with TBS2. The digested peptides in diluted PTS buffer were heated at 95 °C for 10 min to inactivate trypsin, diluted 2-fold with TBS2, and incubated with the ACN-prewashed Tamavidin 2-REV beads for 3 h at 4 °C. During the incubation period, 1 mM biotin solution in TBS3 (50 mM Tris-HCl, pH 7.5, and 500 mM NaCl) was passed through GL-Tip SDB. After washing five times with TBS2, the beads were incubated with 200 μL of TBS3 for 15 min at 37 °C as a mock elution. After removing TBS3, biotinylated peptides were eluted for 15 min at 37 °C twice with 100 μL of the biotin solution that had been passed through GL-Tip SDB. The combined eluates were desalted using GL-Tip SDB, evaporated in a SpeedVac concentrator, and redissolved in 0.1% TFA and 3% ACN.

### LC-MS/MS of BioID Samples

Peptides were analyzed by an EASY-nLC 1200 UHPLC-coupled Orbitrap Fusion Tribrid mass spectrometer (Thermo Fisher Scientific). Aliquots (10 μL) of each sample were loaded by 20 μL of 0.1% formic acid in water at a pressure of 200 bar onto a trap column (100 μm × 2 cm, PepMap nanoViper C18 column, 5 μm, 100 Å; Thermo Fisher Scientific). The following mobile phases were used: solvent A (0.1% formic acid in water) and solvent B (80% ACN and 0.1% formic acid in water). The trapped peptides were eluted from an analytical column (75 μm × 15 cm, nano HPLC capillary column, 3 μm; Nikkyo Technos, Tokyo, Japan) at a constant flow rate of 300 nL/min by changing the gradient: 5% B at 0 min, 40% B at 60 min, 100% B at 70 min, and 100% B at 80 min. The Orbitrap Fusion was operated in positive ion mode with nanospray voltage set at 2.0 kV, source temperature at 275 °C, and S-lens RF level at 60. A scan cycle comprised MS1 scan (*m/z* range from 375 to 1,500) with default charge set to 2, maximum ion injection time of 50 ms, a resolution of 120,000, and automatic gain control (AGC) value of 4e5. The MS1 scan cycle was followed by MS2 scans (ion trap) of the most intense ions fulfilling predefined selection criteria (AGC: 1e5; maximum ion injection time: 200 ms; isolation window: 1.6 *m/z;* spectrum data type: centroid; exclusion of unassigned, singly, and >5 charged precursors; peptide match preferred; exclude isotopes on; dynamic exclusion time: 10 s) for 3 sec. The HCD collision energy was set to 30% normalized collision energy.

### MS Data Analysis of BioID Samples

Raw data were directly analyzed against the Swiss-Prot database, version 2017-10-25, restricted to *Mus musculus* (25,097 sequences) using Proteome Discoverer version 2.4 (Thermo Fisher Scientific) with the Sequest HT search engine. The search parameters were as follows: (a) trypsin as an enzyme with up to two missed cleavages; (b) precursor mass tolerance of 10 ppm; (c) fragment mass tolerance of 0.6 Da; (d) carbamidomethylation of cysteine as a fixed modification; (e) acetylation of protein N terminus, oxidation of methionine, and biotinylation of lysine as variable modifications. Peptides were filtered at a false discovery rate (FDR) of 1% using the Percolator node of Proteome Discoverer.

### TMT-Based Quantitative Proteomics of Surface-Biotinylated Cells

EGF-stimulated and surface-biotinylated HeLa cells were lysed in guanidine buffer as described above. After heating and sonication, proteins (100 μg each) were purified by methanol–chloroform precipitation and resuspended in 20 μL of 0.1% RapiGest SF in 50 mM triethylammonium bicarbonate. After sonication and heating at 95°C for 10 min, the proteins were digested with 1 μg trypsin/Lys-C mix (Promega) at 37 °C overnight. After heating at 95°C for 10 min, the digested peptides from each sample were labeled with 0.1 mg of TMT 11-plex reagents (Thermo Fisher Scientific) and pooled. The combined sample was then diluted 6-fold with TBS2 and incubated with the ACN-prewashed Tamavidin 2-REV beads for 3 h at 4 °C. Biotinylated peptides were enriched by the optimized workflow as described above. LC-MS/MS analysis of the resultant peptides was performed on an EASY-nLC 1200 UHPLC connected to a Q Exactive Plus mass spectrometer through a nanoelectrospray ion source (Thermo Fisher Scientific). The peptides were separated on the analytical column (75 μm × 15 cm, 3 μm; Nikkyo Technos) with a linear gradient of 4–20% for 0–115 min and 20–32% for 115–160 min, followed by an increase to 80% ACN for 10 min and finally hold at 80% ACN for 10 min. The mass spectrometer was operated in data-dependent acquisition mode with a top 10 MS/MS method. MS1 spectra were measured with a resolution of 70,000, an AGC target of 3e6, and a mass range from 375 to 1,400 *m/z*. MS/MS spectra were triggered at a resolution of 35,000, an AGC target of 1e5, an isolation window of 0.7 *m/z*, a maximum injection time of 150 ms, and a normalized collision energy of 34. Dynamic exclusion was set to 20 s. Raw data were directly analyzed against the SwissProt database restricted to *Homo sapiens* using Proteome Discoverer version 2.4 with the Sequest HT search engine for identification and TMT quantification. The search parameters were as follows: (a) trypsin as an enzyme with up to two missed cleavages; (b) precursor mass tolerance of 10 ppm; (c) fragment mass tolerance of 0.02 Da; (d) TMT of peptide N-terminus and carbamidomethylation of cysteine as fixed modifications; (e) TMT of lysine, biotinylation of lysine, and oxidation of methionine as variable modifications. Peptides were filtered at an FDR of 1% using the Percolator node. TMT quantification was performed using the Reporter Ions Quantifier node. Normalization was performed such that the total sum of the abundance values for each TMT channel over all peptides was the same.

### Data Visualization

Results were analyzed using R version 4.1.1 and plotted using RStudio version 1.4.1717. The peptide coverage of Reticulon-4 protein in Figure 4C was illustrated using Protter (27). Proteins with the GO annotation terms “cell surface” and/or “plasma membrane” and/or “extracellular” listed in UniProtKB were classified as cell surface proteins as reported previously (28, 29). The schematic diagram of EGFR in Figure 5E was illustrated using Pfam (30).

**Figure 1.**
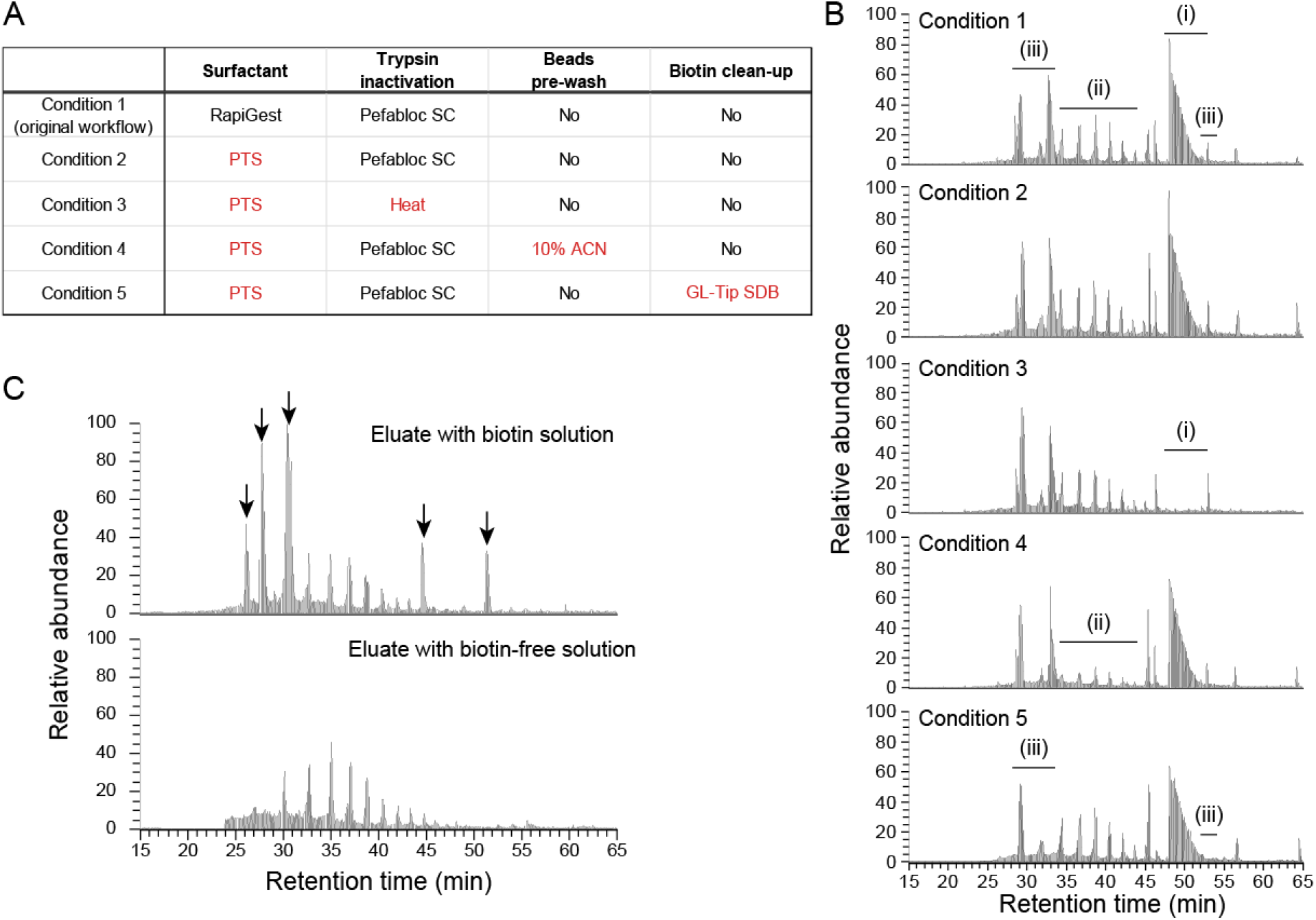
Interference ions generated under various conditions to enrich biotinylated peptides. (A) Table showing the conditions tested to reduce interference ions. Changes from the original workflow are highlighted in red. (B) MS1 chromatograms of peptide samples obtained under each condition. Interference ions indicated by (i), (ii), and (iii) are derived from Pefabloc SC, Tamavidin 2-REV beads, and biotin solution, respectively. Relative abundance is on the same scale across conditions. (C) MS1 chromatograms of the eluate with biotin solution (TBS2 containing 1 mM biotin, top) or with biotin-free solution (TBS2, bottom) from Tamavidin 2-REV beads.

**Figure 2.**
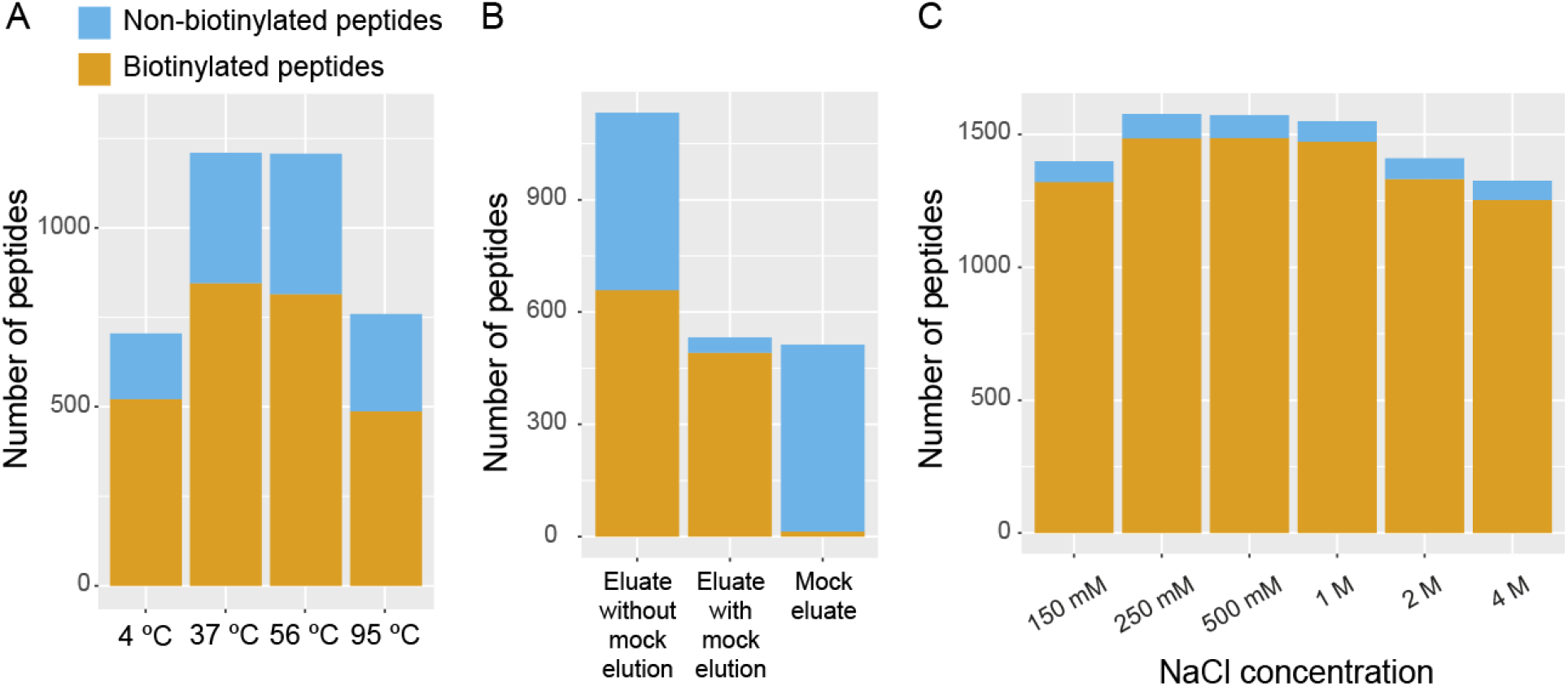
Optimal elution conditions for biotinylated peptides. (A) Comparison of temperatures of biotin solution to elute biotinylated peptides from Tamavidin 2-REV beads. (B) Effect of ‘mock’ elution on the contamination by non-biotinylated peptides in the eluates. (C) Comparison of NaCl concentrations of biotin solution to elute biotinylated peptides.

**Figure 3.**
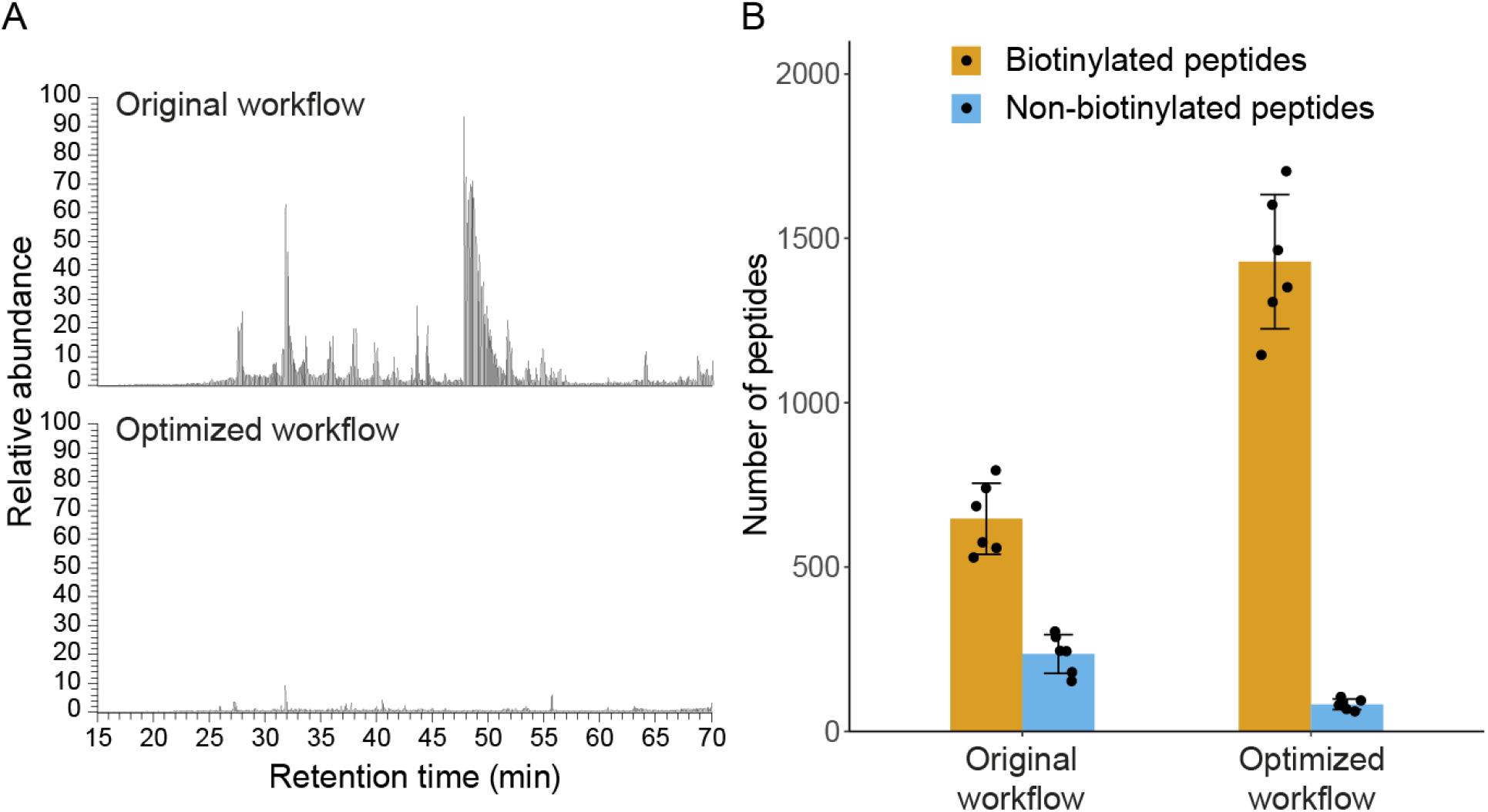
Comparison of biotinylated peptides enriched by the original and optimized workflows. (A) MS1 chromatograms of peptide samples obtained by the original workflow (top) and the optimized workflow (bottom). Relative abundance is on the same scale between the two workflows. (B) The numbers of biotinylated peptides and non-biotinylated peptides that were identified by the original workflow or the optimized workflow.

**Figure 4.**
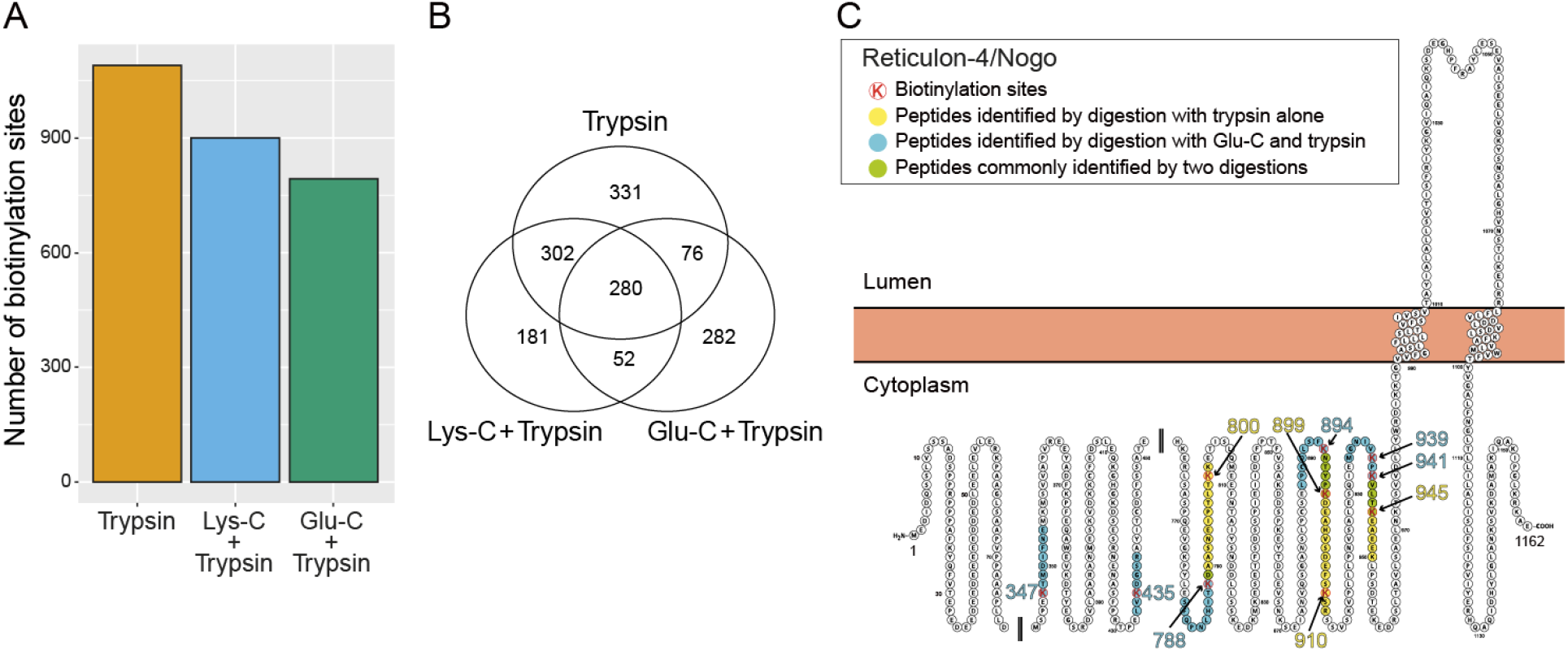
Biotinylation sites identified by digestion with combinations of different proteases. (A) The numbers of biotinylation sites identified by digestion with trypsin alone, Lys-C followed by trypsin, and Glu-C followed by trypsin. (B) Venn diagram showing the distribution of biotinylation sites identified by digestion with the three combinations of different proteases. (C) Schematic of biotinylated peptides and their sites identified by digestion with trypsin alone or Glu-C followed by trypsin on an ER protein Reticulon-4/Nogo.

**Figure 5.**
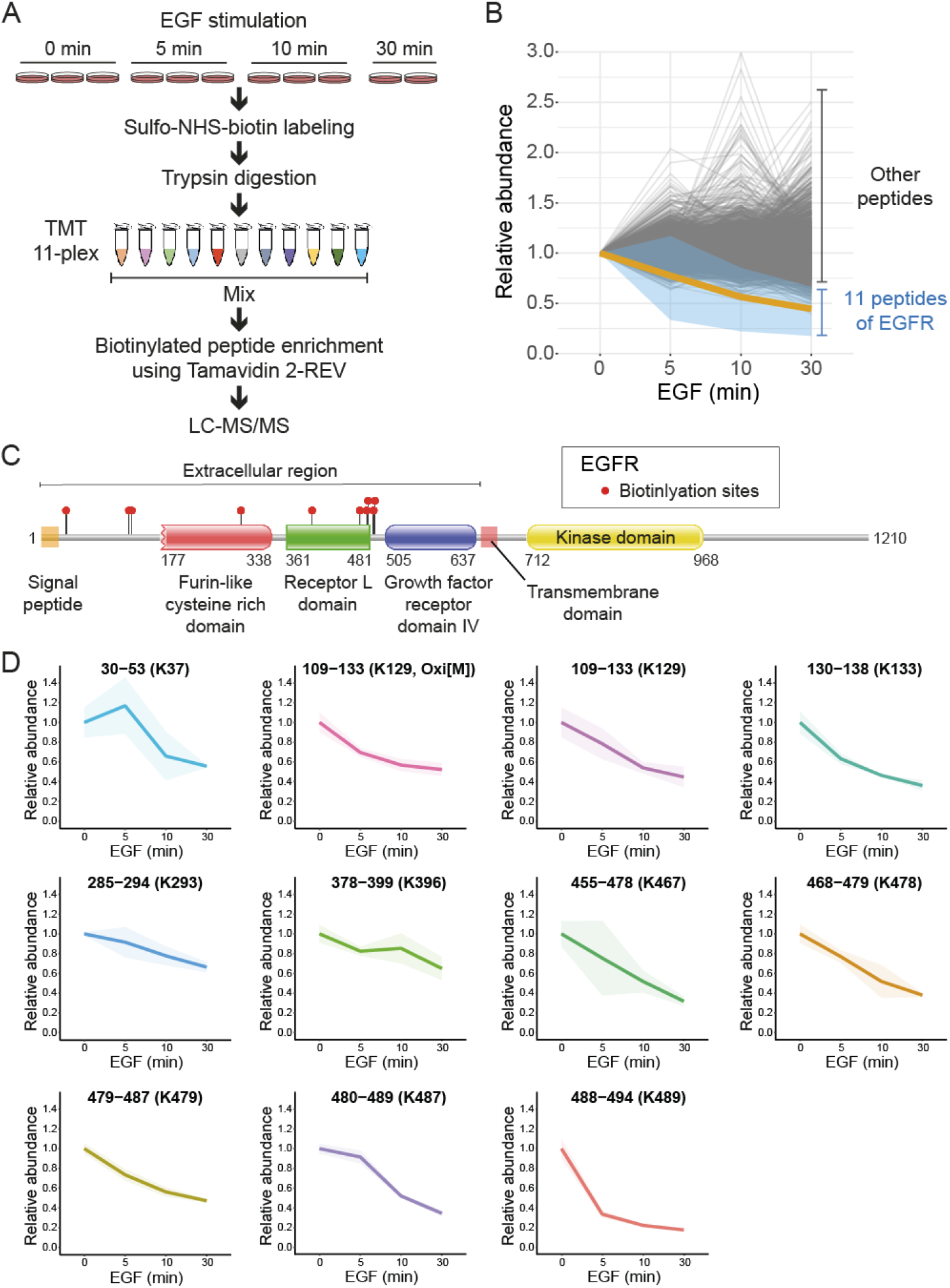
TMT-based multiplexed quantification of biotinylated peptides from EGF-stimulated and surface-biotinylated HeLa cells. (A) Schematic workflow of quantitative cell surface proteomics using surface biotinylation, TMT labeling, and Tamavidin 2-REV enrichment. (B) Changes in relative abundances of all 5,272 biotinylated peptides classified as ‘Surface proteins’ after EGF stimulation. The thick orange line and the surrounding blue ribbon indicate the mean abundance and the area between maximum and minimum abundance of 11 biotinylated peptides derived from the extracellular region of EGFR, respectively. Other biotinylated peptides are shown in gray. (C) Schematic diagram of EGFR with identified biotinylation sites in the extracellular region. (D) EGF-induced decrease of the 11 biotinylated peptides derived from the extracellular region of EGFR. Lines and surrounding ribbons indicate changes in mean abundances and standard deviations between biological replicates, respectively.

### Western Blotting

Cell lysates prepared in guanidine buffer as described above were subjected to methanol–chloroform precipitation and resuspended in SDS-sample buffer. The proteins (10 μg each) were separated by electrophoresis on a 9% polyacrylamide gel and transferred onto a PVDF membrane. After blocking with Blocking One (Nacalai Tesque) for 1 h at room temperature, the membrane was incubated with primary antibodies in TBS-T (20 mM Tris-HCl, pH 7.5, 150 mM NaCl, and 0.05% Tween-20) containing 5% Blocking One overnight at room temperature. After washing twice with TBS-T for 10 min, the membrane was incubated with HRP-linked secondary antibodies for 1 h and then washed twice with TBS-T for 10 min. To detect biotinylated proteins, the transferred membrane was blocked with Blocking One and incubated with HRP-linked streptavidin (BioLegend, #405210, 1:10,000) in TBS-T containing 5% Blocking One for 1 h. Protein bands on the membrane were detected using the ChemiDoc Touch Imaging System (Bio-Rad) after incubation with Clarity Western ECL substrate (Bio-Rad). Antibodies and dilutions used in this study were as follows: anti-STING (D2P2F) rabbit mAb (Cell Signaling Technology, #13647, 1:10,000); anti-EGF receptor (D38B1) XP rabbit mAb (Cell Signaling Technology, #4267, 1:1,000); anti-phospho-ERK1/2 (Thr202/Tyr204) (D13.14.4E) XP rabbit mAb (Cell Signaling Technology, #4370, 1:2,500); anti-α-tubulin (DM1a) mouse mAb (Sigma-Aldrich, T6199, 1:4,000); anti-rabbit IgG, HRP-linked antibody (Cell Signaling Technology, #7074, 1:10,000); anti-mouse IgG, HRP-linked antibody (Cell Signaling Technology, #7076, 1:10,000).

## RESULTS AND DISCUSSION

### Reduction of Interference Ions Generated in the Enrichment Process of Biotinylated Peptides

We have previously established STING-knockout (SKO) RAW264.7 macrophages stably expressing TurboID-fused STING (17). Western blot analysis confirmed the expression of TurboID-STING in these cells and its ability to biotinylate cellular proteins by adding 500 μM biotin to the culture medium for 10 min (Figure S1). This biotinylated sample was used throughout the experiments to optimize the enrichment workflow for biotinylated peptides.

In the original workflow, cellular proteins were extracted with guanidine hydrochloride together with one-step reduction and alkylation, and then purified by methanol–chloroform precipitation. After digestion with trypsin in RapiGest surfactant, Pefabloc SC was added to inhibit trypsin activity. Biotinylated peptides were then captured with Tamavidin 2-REV magnetic beads and eluted competitively with excess biotin (17). When the peptide samples obtained by this workflow were subjected to LC-MS/MS, high intensity peaks of interference and/or contaminant ions were observed at the MS1 level, particularly when small amounts of biotinylated peptides were analyzed (Figure 1 A,B, Condition 1). These peaks could cause ion suppression and inefficient identification of biotinylated peptides. Therefore, we attempted to optimize each step of the enrichment process to reduce these interference ions and efficiently elute biotinylated peptides from Tamavidin 2-REV.

First, to examine the effect of surfactants that solubilize precipitated proteins and promote enzymatic digestion (31–33), we compared RapiGest (34) with a phase-transfer surfactant (PTS) composed of sodium deoxycholate (SDC) and sodium lauroylsarcosinate (SLS) (24–26). As a result, there were almost no differences in MS1 chromatograms between RapiGest and PTS (Figure 1A,B, Condition 1 vs. Condition 2); however, we started using PTS because it is more cost-effective than RapiGest.

Second, we focused on the serine protease inhibitor Pefabloc SC, 4-(2-aminoethyl)-benzenesulfonyl fluoride (35) that was used to inactivate trypsin and prevent digestion of Tamavidin 2-REV when capturing biotinylated peptides. Because trypsin is inactivated by heat (36, 37), we compared the effects of Pefabloc SC and heat treatment on MS1 chromatograms and found that the interference ions observed around 50 min (Figure 1B, (i)) were dramatically reduced in the heat-treated sample (Figure 1A,B, Condition 2 vs. Condition 3). These results indicate that heat treatment is superior to the addition of Pefabloc SC.

Third, we examined a series of interference ions detected between 35 min and 45 min on MS1 chromatograms (Figure 1B, (ii)). The repeating and characteristic *m/z* patterns suggested contamination by polymer-like materials from the Tamavidin 2-REV magnetic beads. Therefore, when the beads were pre-washed with 10% acetonitrile before capturing biotinylated peptides, these interference ions were clearly reduced (Figure 1A,B, Condition 2 vs. Condition 4).

Fourth, to investigate contamination in the biotin solution used to elute biotinylated peptides from the beads, we compared eluates with the biotin solutions with or without passing through the GL-Tip SDB desalting tip. The results showed that some contaminant ions (Figure 1B, (iii)) were removed when the biotin solution had been passed through GL-Tip SDB (Figure 1A,B, Condition 2 vs. Condition 5). To confirm that these contaminant ions were not derived from digested peptides but from the biotin solution, Tamavidin 2-REV beads that had not been incubated with the digested peptides were eluted with the biotin solution or biotin-free solution, and the eluates were analyzed by LC-MS/MS. While five intense peaks were detected in the eluate with the biotin solution (Figure 1C, arrows), these peaks were strongly reduced in the eluate with the biotin-free solution. These contaminants had hydrophobic properties and could not be removed by desalting the eluates using GL-Tip SDB. Thus, we eluted biotinylated peptides with the biotin solution that had been passed through GL-Tip SDB.

### Optimization of Elution Conditions for Biotinylated Peptides from Tamavidin 2-REV

Next, we attempted to optimize the conditions for competitive elution of biotinylated peptides from the Tamavidin 2-REV beads. First, we focused on the temperature at which biotinylated peptides were eluted from the beads. In the original workflow, biotinylated peptides were eluted with the biotin solution at 95 °C. We therefore compared four temperatures: 4, 37, 56, and 95 °C. Among them, the highest number of biotinylated peptides was identified at 37 °C (Figure 2A). The interference ions were increased with the increase in the elution temperature, suggesting that these interference ions were non-specifically eluted from the beads by heating (Figure S2).

We next investigated whether the contamination by non-biotinylated peptides in the eluates could be reduced by adding a step of ‘mock’ elution with biotin-free buffer immediately before elution with the biotin solution. After the beads were incubated with biotin-free buffer for 15 min at 37 °C, the buffer was removed and biotinylated peptides were eluted from the beads with the biotin solution at 37 °C. As a result, non-biotinylated peptides were efficiently removed from the eluate, which increased the ratio of biotinylated to non-biotinylated peptides (Figure 2B).

The previous crystal structure analysis had revealed that Tamavidin 2 forms multiple hydrogen bonds with biotin (38), and therefore we considered the possibility that biotinylated peptides could be efficiently eluted by increasing the salt concentration to weaken the hydrogen bonds. After the mock elution, biotinylated peptides were eluted from the beads at 37 °C with the biotin solutions containing various concentrations of NaCl. As a result, the highest number of biotinylated peptides was identified around 500 mM NaCl (Figure 2C). On the basis of these results of optimizing the elution conditions, we eluted biotinylated peptides from Tamavidin 2-REV beads with the biotin solution containing 500 mM NaCl at 37 °C after the mock elution.

### Comparison of Biotinylated Peptides Enriched by the Original and Optimized Workflows

To evaluate the optimized workflow, biotinylated peptides enriched by the original and optimized workflows from 300 μg proteins of RAW264.7 cells expressing TurboID-STING were compared by LC-MS/MS analysis. MS1 chromatograms showed that many peaks of interference ions in the original workflow were greatly reduced in the optimized workflow (Figure 3A). Furthermore, the number of identified biotinylated peptides was increased by more than two-fold after the optimization, and the contamination by non-biotinylated peptides was reduced to almost one-third, resulting in approximately 95% of all identified peptides being biotinylated (Figure 3B). Thus, we have established an optimized workflow to efficiently and specifically enrich biotinylated peptides using Tamavidin 2-REV.

### Sequential Digestion with Different Proteases

BioID-based proximity biotinylation of proteins occurs on lysine residues, which can inhibit efficient digestion with trypsin (39, 40). Previous studies of cross-linking MS using lysine-reactive cross-linkers (*e.g.*, DSSO) have shown that sequential digestion with Lys-C or other non-lysine-specific proteases (*e.g.*, Glu-C) followed by trypsin increases the coverage of identified cross-linked peptides (41, 42). We therefore examined three combinations of different proteases to digest biotinylated proteins from TurboID-STING-expressing cells: trypsin alone, Lys-C followed by trypsin, and Glu-C followed by trypsin. First, we examined whether PTS buffer used to solubilize precipitated proteins was compatible with digestion with these proteases, and confirmed that each protease cleaved at specific residues; that is, > 97% of all cleavages by trypsin occurred at lysine or arginine residues, >96% of all cleavages by Lys-C occurred at lysine residues, and 97% of all cleavages by Glu-C occurred at aspartate or glutamate residues (Figure S3) (43). Next, when biotinylated peptides were enriched after digestion with the three combinations of different proteases, digestion with trypsin alone identified the highest number of biotinylation sites (Figure 4A) and the highest number of unique biotinylation sites (Figure 4B). Sequential digestion with Lyc-C and trypsin yielded similar results to digestion with trypsin alone. For example, many of identified biotinylation sites were shared by the two digestions (Figure 4A,B). On the other hand, sequential digestion with Glu-C and trypsin yielded many unique biotinylation sites that were not identified by trypsin alone or by Lys-C and trypsin, although the total number of identified biotinylation sites was less (Figure 4A,B).

Consistent with the endoplasmic reticulum (ER) localization of STING at the steady state, many biotinylation sites identified by the three digestions were located on ER proteins (Table S1). In the case of an ER protein Reticulon-4/Nogo (44, 45), while four biotinylation sites were identified by digestion with trypsin alone, six other biotinylation sites were identified by sequential digestion with Glu-C and trypsin (Figure 4C). All 10 biotinylation sites were located in the N-terminal cytoplasmic region of Reticulon-4, consistent with the fusion of TurboID to the cytoplasmic side of STING. Collectively, these results indicate that digestion with trypsin alone is most effective in identifying peptides with biotinylated lysine residues. In addition, sequential digestion with Glu-C and trypsin can identify further biotinylation sites and provide more detailed topological information of membrane proteins.

### Multiplexed Quantitative Cell Surface Proteomics by the Combination of TMT Labeling and Biotinylated Peptide Enrichment

Finally, quantitative cell surface proteome profiling was performed by applying this optimized workflow. Selective biotinylation of surface-exposed primary amines on cell surface proteins is one of the most frequent approaches, and the membrane-impermeable reagent Sulfo-NHS-biotin is typically used (40). We examined changes in the surface proteome of HeLa cells stimulated with epidermal growth factor (EGF), which is well known to induce the internalization and degradation of EGF receptor (EGFR) (46). First, we confirmed that EGF stimulation of HeLa cells induced rapid and transient phosphorylation of ERK1/2 and gradual degradation of EGFR, as previously reported (Figure S4A) (47, 48). Next, surface biotinylation was performed at four time points after EGF stimulation in multiple biological replicates, followed by digestion of cellular proteins with trypsin. After TMT labeling and mixing of digested peptides, biotinylated peptides were enriched using Tamavidin 2-REV (Figure 5A). In total, 7,996 biotinylated peptides were identified and quantified by LC-MS/MS analysis (Table S2) and classified as peptides of ‘Surface proteins’ or ‘Non-surface proteins’ according to the GO cellular component annotation used in previous reports (28, 29). The number of quantified biotinylated peptides classified as ‘Surface proteins’ accounted for 66% of the total (Figure S4B), indicating that biotinylated peptides derived from cell surface proteins were efficiently enriched. We further examined changes in relative abundances of the biotinylated peptides classified as ‘Surface proteins’ during the time course of EGF stimulation (Figure 5B). As a result, 11 biotinylated peptides derived from the extracellular region of EGFR were detected (Figure 5C,D), and all 11 biotinylated peptides were specifically decreased by EGF stimulation (Figure 5B,D), confirming the specific internalization of EGFR by EGF stimulation. These results clearly demonstrated that the combination of TMT-based multiplexing and this optimized workflow enables large-scale quantification of the cell surface proteome.

## CONCLUSIONS

Global identification of protein–protein interactions and subcellular proteomes in living cells is essential to elucidate regulatory mechanisms of various biological processes. Proximity-dependent biotinylation has emerged as a promising approach that overcomes the limitations of classical biochemical fractionation and affinity purification. Various techniques for proximity-dependent biotinylation followed by MS analysis have been rapidly developed and improved. In particular, BioID enzymes have been subjected to intensive engineering and optimization. Although specific enrichment of biotinylated proteins/peptides from complex biological samples is a critical step for proximity-dependent identification, the enrichment methods have not been fully developed. In this study, we have established an optimized workflow for enrichment and identification of biotinylated peptides using Tamavidin 2-REV. In contrast to the conventional enrichment and identification of biotinylated proteins using streptavidin, identification of biotinylated peptides provides information about the biotinylation sites on proteins. In addition to trypsin digestion alone, sequential digestion with Glu-C and trypsin can provide more comprehensive biotinylation sites and more detailed topological information of membrane proteins. Because advances in proximity-dependent biotinylation and chemical biology such as acyl-biotin exchange are increasing the importance of MS-based identification of biotinylated proteins and their sites, this optimized workflow may be useful for a wide range of applications.

## Author Contributions

K.N., H.Y., and H.K. conceived the project and analyzed the data. K.N., H.Y., and K.M. performed the experiments. H.Y. and H.K. wrote the manuscript. H.K. supervised the project. All authors read and approved the final manuscript.

## Notes

The authors declare no competing financial interest. The mass spectrometry raw files and search results have been deposited in the ProteomeXchange Consortium via the jPOST partner repository (49) with the dataset identifiers PXD032059, PXD032060, PXD032061, and PXD032062.

## ACKNOWLEDGMENTS

This work was supported by the JSPS KAKENHI (17K08661 and 20K07340 to K.M., 21K06138 to H.Y., and 19H04966 and 20K06628 to H.K.) and the Takeda Science Foundation (to K.M. and H.Y.). We thank Megumi Kawano and Mayumi Kajimoto for technical assistance and Mayumi Iwata for secretarial assistance. We also thank Mitchell Arico from Edanz (https://jp.edanz.com/ac) for editing a draft of this manuscript.

