## Supplementary Figures for "Optimized Workflow for Enrichment and Identification of Biotinylated Peptides using Tamavidin 2-REV for BioID and Cell Surface Proteomics"

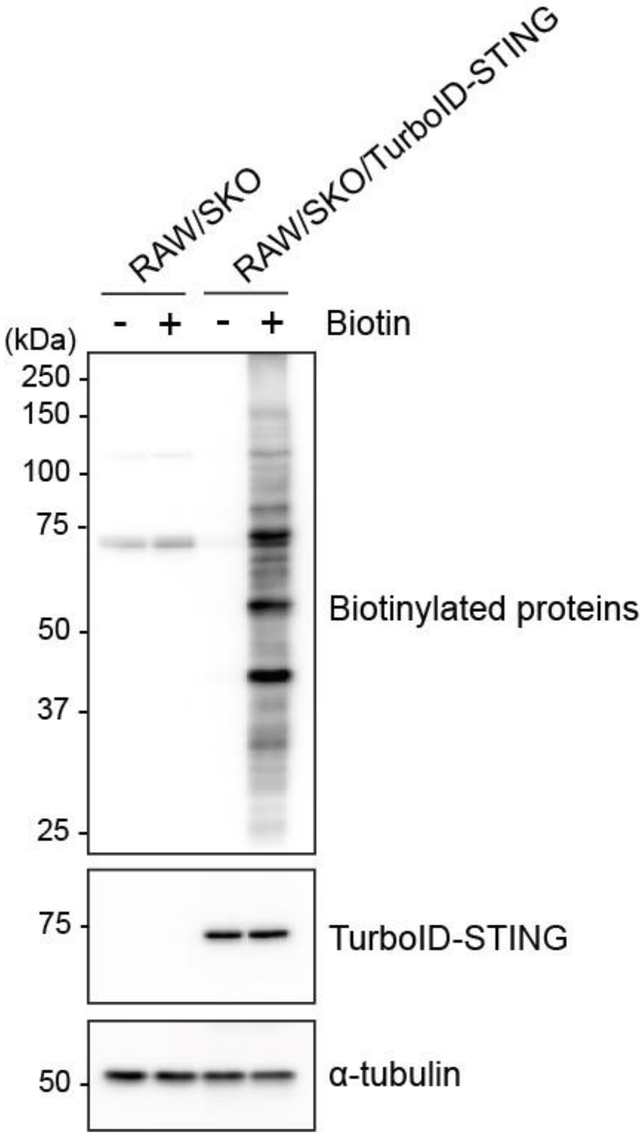


**Figure S1.** Biotinylation of cellular proteins by TurboID-STING.

STING-knockout (SKO) RAW264.7 cells and SKO RAW264.7 cells expressing TurboID-STING were cultured for 10 min in the presence or absence of 500 μM biotin. The cell lysates were analyzed by Western blotting with HRP-conjugated streptavidin and antibodies to STING and α-tubulin.

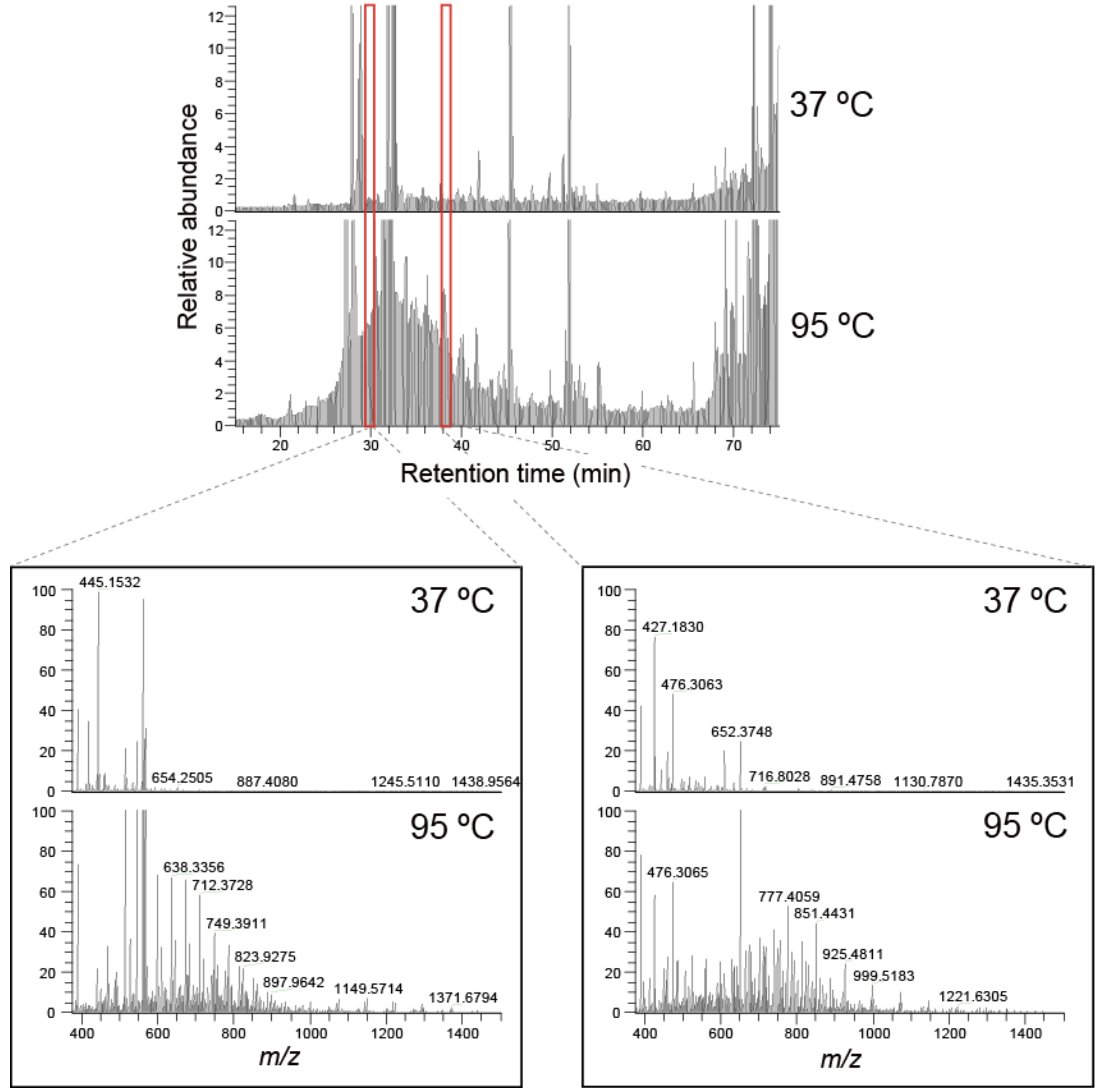


**Figure S2.** Interference ions eluted by heating Tamavidin 2-REV beads.

MS1 chromatograms of eluates from Tamavidin 2-REV beads at 37 °C and 95 °C (top). Bottom shows interference ions eluted at 95 °C during the time windows indicated by red boxes.

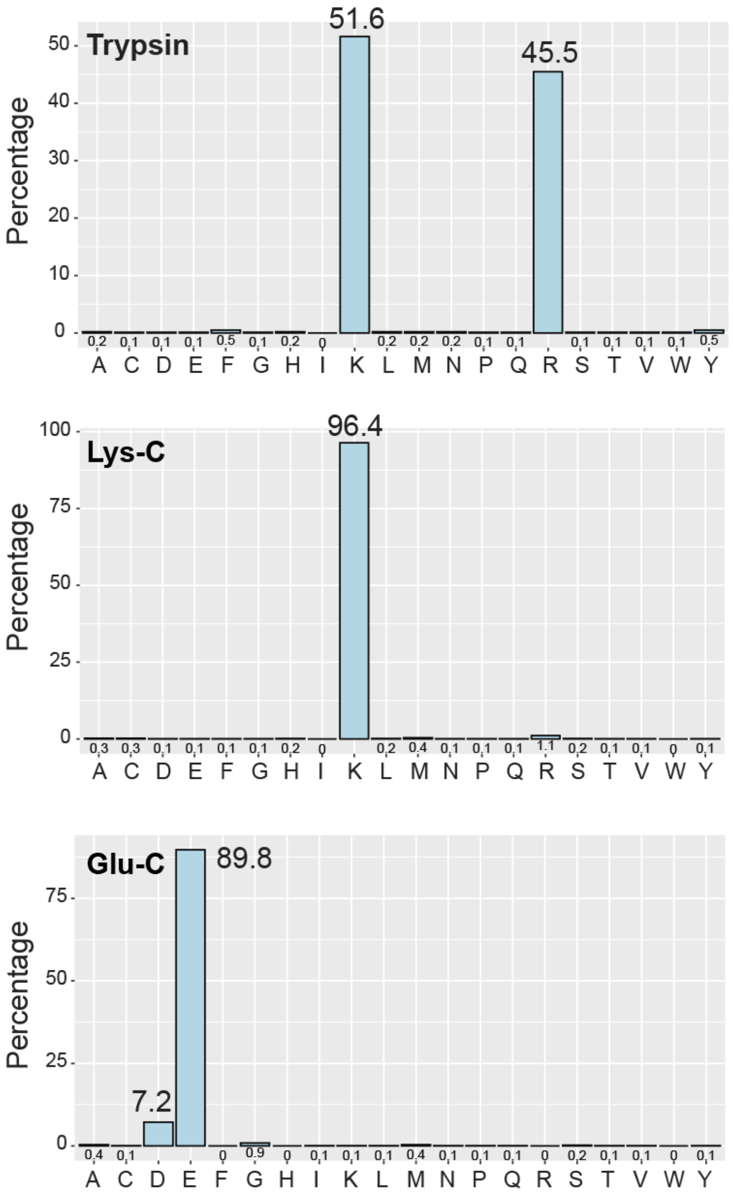


**Figure S3.** Digestion specificity of trypsin, Lys-C, and Glu-C in PTS buffer.

Proteins (60 µg) were purified by methanol–chloroform precipitation and resuspended in 20 µL of PTS buffer (100 mM Tris-HCl, pH 8.0, 12 mM SDC, and 12 mM SLS). After 5-fold dilution with 100 mM Tris-HCl, pH8.0, the protein solution was divided into three equal portions and digested with 200 ng each of trypsin, Lys-C, or Glu-C at 37 °C overnight. The percentages of the last amino acid residues of identified peptides were calculated and plotted.

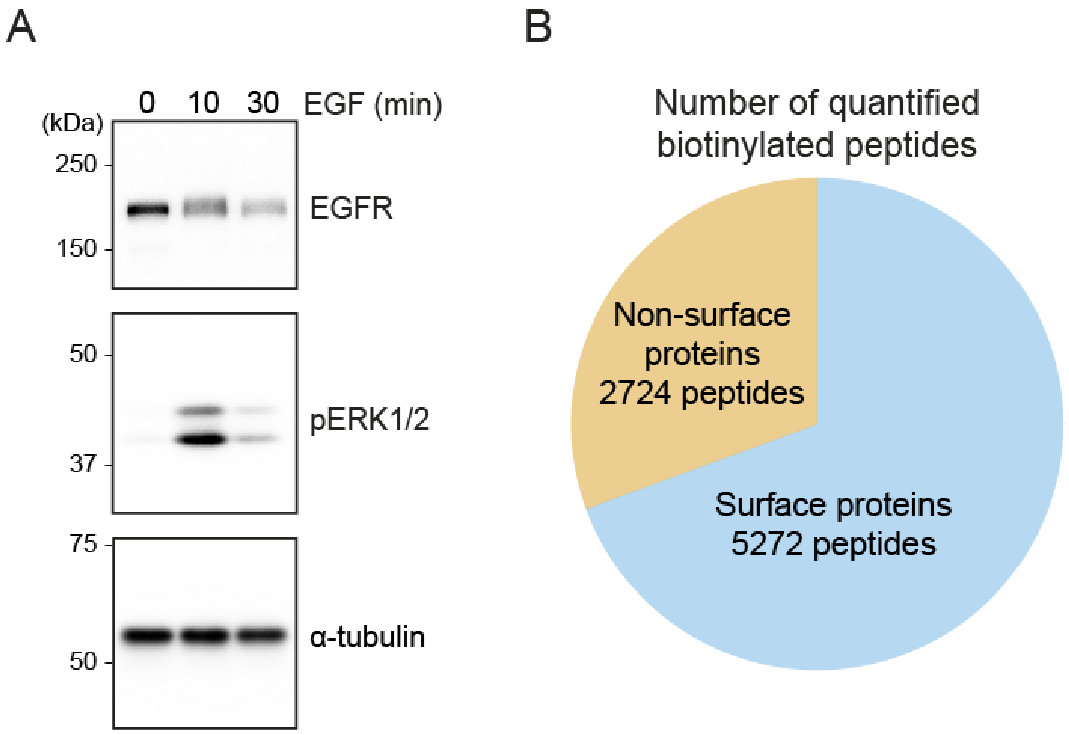


**Figure S4.** EGF stimulation and surface biotinylation of HeLa cells.

(A) HeLa cells were starved in serum-free DMEM for 18 h and then stimulated with 100 ng/mL EGF for 0, 10, or 30 min. The cell lysates were analyzed by Western blotting with antibodies to EGFR, phosphorylated ERK1/2, and α-tubulin. (B) Pie chart showing the number of quantified biotinylated peptides classified as ‘Surface proteins’ or ‘Non-surface proteins’.
